# Human trophoblast stem cells can be used to model placental susceptibility to *Toxoplasma gondii* and highlight the critical importance of the trophoblast cell surface in pathogen resistance

**DOI:** 10.1101/2023.11.10.566663

**Authors:** Rafaela J. da Silva, Leah F. Cabo, Jada L. George, Laty A. Cahoon, Liheng Yang, Carolyn B. Coyne, Jon P. Boyle

## Abstract

The placenta is a critical barrier against viral, bacterial, and eukaryotic pathogens. For most teratogenic pathogens, the precise molecular mechanisms of placental resistance are still being unraveled. Given the importance to understand these mechanisms and challenges in replicating trophoblast-*pathogen* interactions using *in vitro* models, we tested an existing stem-cell derived model of trophoblast development for its relevance to infection with *Toxoplasma gondii*. We grew human trophoblast stem cells (TS^CT^) under conditions leading to either syncytiotrophoblast (TS^SYN^) or cytotrophoblast (TS^CYT^) and infected them with *T. gondii*. We evaluated *T. gondii* proliferation and invasion, cell ultrastructure, as well as for transcriptome changes after infection. TS^SYNs^ cells showed similar ultrastructure compared to primary cells and villous explants when analyzed by TEM and SEM, a resistance to *T. gondii* adhesion could be visualized on the SEM level. Furthermore, TS^SYNs^ were highly refractory to parasite adhesion and replication, while TS^CYT^ were not. RNA-seq data on mock-treated and infected cells identified differences between cell types as well as how they responded to *T. gondii* infection. We also evaluated if TS^SC^-derived SYNs and CYTs had distinct resistance profiles to another vertically transmitted facultative intracellular pathogen, *Listeria monocytogenes*. We demonstrate that TS^SYNs^ are highly resistant to *L. monocytogenes*, while TS^CYTs^ are not. Like *T. gondii*, TS^SYN^ resistance to *L. monocytogenes* was at the level of bacterial adhesion. Altogether, our data indicate that stem-cell derived trophoblasts recapitulate resistance profiles of primary cells to *T. gondii* and highlight the critical importance of the placental surface in cell-autonomous resistance to teratogens.

## INTRODUCTION

Congenital infection occurs when a fetus contracts an infection from the mother during pregnancy. The impact on the developing fetus can vary depending upon factors such as the gestational age during the infection and the specific pathogen responsible, resulting in a wide array of outcomes including miscarriage, stillbirth, fetal malformation, and neonatal death. Pathogens such as *Toxoplasma gondii* and *Listeria monocytogenes* are important among the major causes of congenital infections and related to several adverse fetal and neonatal outcomes (1, 2).

*Toxoplasma gondii* is an obligatory intracellular protozoan parasite responsible for the clinical illness toxoplasmosis and is particularly important as a causative agent of disease in the immunocompromised and pregnant individuals (3, 4). In immunocompetent patients, toxoplasmosis is generally asymptomatic (5). However, congenital toxoplasmosis, whereby an immunocompetent mother transmits the parasite to their developing fetus, can be lethal (6, 7). Congenital toxoplasmosis is one of the most severe forms of the disease with primary infection during pregnancy resulting in miscarriage, stillbirth, premature birth, malformations, and neurological and/or ocular disorders in newborns (4, 8–10).

To reach the fetus and cause congenital toxoplasmosis, *T. gondii* must cross the barriers protecting the fetus, including the placenta (6, 7, 11). This organ is the primary site of nutrient and gas exchange between mother and fetus and *T. gondii* is capable of broad dissemination within the host via the bloodstream, highlighting the importance of encounters between *T. gondii* and the placenta. The placenta also produces hormones and functions as an immunological and physical barrier to bloodborne pathogens (2, 12–14). *T. gondii* infection of the fetus is not the rule and occurs approximately in 40% of pregnant women who are infected for the first-time during gestation (15). It is likely, but yet unproven, that the placenta protects the fetus from infection in at least some of these cases.

Structurally, the placenta is formed by villous trees that are either free-floating and bathed in maternal blood or anchored in the decidua. The inner layer of each villous tree is composed of cytotrophoblast (CYT). CYT are mononucleated cells that are responsible for a) replenishing and growing the protective syncytiotrophoblast (SYN) layer via cell fusion and b) differentiating into extravillous trophoblast cells (EVT) (2, 16). The SYN layer is made up of a multinucleated cell that is bathed in maternal blood and present on the outermost surface of floating villous (2, 17, 18). In contrast, EVTs are mononucleated, mesenchymal cells with an invasive profile that anchor the placenta in the decidua, where they then interface with maternal decidual and immune cells (19, 20). The SYN layer is a critical component of fetal defense and in recent years has been found to be naturally pathogen resistant, including to viral pathogens like Zika virus (21),bacterial pathogens like *L. monocytogenes* (22, 23) and parasites like *T. gondii* (18, 24). Our prior work with *T. gondii* using primary human trophoblasts (18) has shown that SYNs resist *T. gondii* infection by a) being refractory to parasite adhesion and b) restricting parasite replication and/or being parasiticidal (18, 24, 25). In contrast, and like nearly all other cell types studied to date, CYTs and EVTs are both susceptible to *T. gondii* infection (24).

The intrinsic mechanisms involved in restricting pathogen growth and invasion by SYN and mechanisms related to susceptibility of CYT cells to the parasite are poorly understood. For *T. gondii*, SYNs represent one of the only known cell types that resist *T. gondii* adhesion and restrict its replication without treatment with interferon-γ, as this parasite is capable of infecting and thriving within most nucleated cells. *In vitro* models that faithfully replicate CYT, SYN and EVT biology are critical for understanding these processes on the molecular level. While lineages of immortalized trophoblast cells derived from choriocarcinomas are often used, including BeWo, JEG-3, and JAR, even when they are syncytialized they do not reproduce the sensitivity of primary trophoblast cells or villous explants (18, 25). While primary trophoblast cells can differentiate spontaneously into SYN, they present challenges of their own, in that they are difficult to manipulate genetically due to their short lifespan *in vitro* (26).

In order explore different cellular models to study the placenta cells and pathogens interactions, we were interested in characterizing and utilizing the human trophoblast stem cells (TS^CT^), previously isolated and described by Okae and collaborators (27) as a model a model to elucidate the differential susceptibility to *Toxoplasma* infection. Here we investigate the utility of TS cells to study the genetics of resistance and susceptibility between SYN and CYT to *T. gondii* and find that they faithfully recapitulate the resistance profile of primary cells to both *T. gondii* and *Listeria monocytogenes*. We also find that they have important limitations with respect to their constitutive production of cytokines like interferon-λ and ability to respond to *T. gondii* infection by the production of CCL22 (18, 28).

## MATERIALS AND METHODS

### Culture of human trophoblast stem cells (TS^CT^)

Trophoblast stem cells (TS^CT^) (clone 27), derived from first trimester placental tissue were kindly provided by Professor Okae from the Tohoku University, Japan. Cells from this line were cultured as described previously (27). Briefly, 75 cm^2^ flasks were incubated in TS medium containing 2 μg/mL iMatrix-55 (AMSBIO, Abingdon, UK) for 10 minutes at 37°C in 5% CO_2_. TS medium consist of basal medium (DMEM/F12 (Gibco, Waltham, MA, USA), 1% ITS-X100 (ThermoFisher Scientific, Waltham, MA, US), 0.3% acid fatty free BSA (Sigma, St Louis, MO, USA), 200 μM of ascorbic acid (Sigma), and 0.5% Penicillin-Streptomycin (ThermoFisher Scientific) and 0.5% of KSR (Gibco), supplemented with 25 ng/EGF (ThermoFisher Scientific), 2 μM CHIR99021 (Stemolecule Reprocell USA, Inc, Beltsville, MD, USA), 5 μM A83-01 (Stemolecule Reprocell USA, Inc, Beltsville, MD, USA), 0.8 mM VPA (APExBIO, Houston, TX, USA) and 5 μM Y27632 (Stemolecule Reprocell USA, Inc, Beltsville, MD, USA). Later, cells were seeded in a ratio of 1:3 and incubated at 37°C and 5% CO_2_, until they reached 80% confluency. After that, cells were collected using TrypLE (Sigma) for 10 min at 37°C and passage to a new pre-coated flask.

### Differentiation of cytotrophoblast cells (TS^CYT^) into syncytiotrophoblast cells (TS^SYN^) and *T. gondii* infection

To induce syncytiotrophoblast (TS^SYN^) development from TS^CYT^ cells, we used both differentiation protocols as outlined in (27), with minor modifications. Briefly, for a mixed population of CYTs and SYNs, TS basal medium was supplemented with 5 μM Y27632 (Stemolecule Reprocell USA, Inc), while for pure populations of TS^SYN^ we used ST (3D) medium [DMEM/F12 (Gibco), ITS-X100 (ThermoFisher Scientific), 0.3% fatty acid free BSA (Sigma), 0.5% Penicillin-Streptomycin (ThermoFisher Scientific), 0.1 mM 2-mercaptoethanol (Fisher Scientific)), supplemented with 2.5 μM Y27632 (Stemolecule Reprocell USA), 2 μM Forskolin (Sigma), 4% EGF (ThermoFisher Scientific), and 50 ng/mL KSR (Gibco)]. In both cases, the medium was added in 6-well plates for 10 min at 37°C, and cells were seeded in a ratio of 1.5 x10^5^ in 6-well plates. For mixed populations, the medium was replaced on day 3. For TS^SYN^ (3D) culture, each well was supplemented with ∼2 mL of ST (3D) media on day 3, and the media was replaced on day 5 with ST (3D) media lacking forskolin for *T. gondii* infection.

### Toxoplasma gondii culture

*Toxoplasma gondii* strain RH (expressing YFP off of the GRA1 promoter) (18) was cultured in human foreskin fibroblast (HFF) in complete Dulbeccòs modified Eagle’s medium (cDMEM; ThermoFisher Scientific plus 100 U/ml penicillin/streptomycin, 100 μg/ml streptomycin, 2 mM L-glutamine, 10% FBS, 3.7 g NaH2CO3/L) and incubated in 5% CO_2_ and 37°C. For infection assays, infected monolayers were scraped and syringe-lysed to release tachyzoites, and then, parasites were pelleted at 800 x *g* for 10 min. For mock infections, the same parasite preparations were passed through a 0.22 µm filter (Millipore, Burlington, MA, US) and the eluate was used to treat host cells at the same dilution as the parasite preparation.

### Listeria monocytogenes strains and culture

We used the following *L. monocytogenes* strains: strain 10403S served as the wild type strain along with isogenic Δ*prsA2* (NF-L1651) (29) Δ*hly* (DP-L2161) (29) strains with deletions in the *prsA2* and *hly* genes, respectively. We also use *Lm*-GFP (DP-L4092) for imaging L. monocytogenes interactions with placental cells (30). For invasion assays, bacteria were grown overnight without agitation at 37°C in brain heart infusion broth (BHI; Oxoid) until an OD_600_ of 0.7-0.9 was reached. Bacteria were washed twice and suspended in cell culture medium without serum and antibiotics for infection experiments.

### General *T. gondii* infection and antibody staining protocol

Trophoblast stem cells grown under different conditions described above were seeded in a ratio of 1.5 x10^5^ in 6-well plate, infected or mock infected with *T. gondii* RH: YFP. TS^CYT^ were induced to form TS^SYN^ for 4-5 days and then infected with *T. gondii* at different MOIs depending on the experiment. Cells were collected and evaluated for 24 and 48h post-infection. For most of 24h infections, TS^SYNs^ were infected on day 5 of differentiation at an MOI of 5 parasites. In both cases the TS^CYT^ were infected 1 day after plating, and the mixed population of trophoblasts were infected in day 3 of differentiation. Cell passages were staggered so that the same parasite preparation could be used to infect TS^SYN^ and TS^CYT^ simultaneously.

To evaluate on which day of differentiation TS^SYN^ becomes resistant to *T. gondii* infection, TS^SYN^ cells were plated in 6-well plated in duplicate and infected with the parasite every day from day 1 to day 6. The parasites were allowed to grow for 24h for each time point and then, the cells were fixed and processed for immunofluorescence.

For immunofluorescence assay, cells were fixed using 4% paraformaldehyde (PFA) (ThermoFisher Scientific) for 12 min, rinsed with PBS (ThermoFisher Scientific) and permeabilized with 0.2% Triton X-100 in PBS for 10 min. Cells were then incubated with the TS^SYN^ marker anti-Syndecan-1 (SDC-1) (1:500) (ab128936, Abcam, Cambridge, UK) and TS^CYT^ marker, anti-Integrin alfa-6 (ITGA6) (1:1000) (MA5-16884, ThermoFisher Scientific) for 1 h at room temperature. Alexa Fluor 594 and 647-conjugated (A-21209 and A-32733, Life Technologies Alexa Fluor H+L, Carlsbad, CA, USA) were used as secondary antibodies for 45 min.

### *Listeria monocytogenes* infection and gentamicin survival assay

TS^SYNs^ and TS^CYTs^ were cultured in 6-well plates and *L. monocytogenes* wildtype, Δ*prsA2* or Δ*hly* was used to infect cells in triplicate at 8 x 10^6^ bacteria per well. As a control, a mock infection was performed using 0.2 μm-filtered *L. monocytogenes* culture supernatant. Inoculated cells were incubated for 1 hour at 37°C with 5% CO_2_. Cells were then washed with fresh cell culture media and incubated for 7 hours with gentamicin (5 µg/ml). Two washes were performed with 1× PBS and cells were lysed by incubation with 0.25% Triton X-100 for 5 min at RT followed by plating of serial dilutions of the cell lysates on BHI agar plates. Plates were incubated at 37°C for 36 hours after which bacteria were enumerated and colony forming units were calculated. TS^SYN^ and TS^CYT^ cells were also collected for immunofluorescence assay as described above and for TEM.

#### RT-qPCR

To quantify the number of parasites in TS^SYN^ and TS^CYT^, cells were infected with *T. gondii* at an MOI of 5 parasites for 24h. After that, genomic DNA was extracted using the GeneJet Genomic DNA Purification kit following manufacturer’s instructions (TermoFisher Scientific). qPCR was used to quantify the total number of genomes using primers targeting the *T. gondii* GRA1 gene and primers targeting human β-ACTIN as a control gene. All reactions were performed in triplicate using a QuantStudio 3 Real-Time PCR System (ThermoFisher). The DNA was mixed with SYBR Green buffer (BioRad, Hercules, CA, USA) and 1 μL (5 µM) of both forward and reverse primers and ddH_2_O. Genes were amplified using a standard protocol (95°C for 10 min and 40 cycles of 95°C for 15 sec and 60°C for 1 min) and data was analyzed with QuantStudio^TM^ Design & Analysis Software. To determine the total number of parasite genomes, a standard curve of known parasite numbers ranging from 1×10^7^ to 1×10^1^ was also performed using *T. gondii* GRA1 primers.

*T. gondii* GRA1 forward: TTAACGTGGAGGAGGTGATTG; GRA1 reverse: TCCTCTACTGTTTCGCCTTTG, and human β-ACTIN forward: GCGAGAAGATGACCCAGATC; human β-ACTIN reverse: CCAGTGGTACGGCCAGAGG. Two experiments in triplicate were performed.

To quantify the expression level of IL-1β in cells infected and mock infected with *L. monocytogenes,* and the RNA were extracted using the Qiagen RNA extraction kit following manufacturer’s instructions (Qiagen, Hilden, Germany). RNA was analyzed by gel electrophoresis and quantified by Nanodrop. cDNA was generated from 0.5 µg of RNA using the SuperScript IV First-Strand synthesis system (ThermoFisher Scientific). IL-1β forward: CTCTCACCTCTCCTACTCACTT; IL-1β reverse: TCAGAATGTGGGAGCGAATG. And β-ACTIN was used as a reference gene. DeltaCt values (query gene Ct – control gene Ct) were used for statistical comparisons and then converted to fold-difference using the 2^-ΔΔCt^ method. One experiment in triplicate was performed.

### Transcriptional analysis of TS^CYT^ and TS^SYN^ using RNAseq

TS^SYN^ and TS^CYT^ cells were seeded and infected in the same conditions as described above for qPCR, and RNA was extracted to perform RNAseq. Strand-specific, oligo-dT generated sequencing libraries were prepared at Core Facility at the University of Pittsburgh and 2x 66 bp reads were sequenced on a NextSeq 200 (Illumina). Read libraries were mapped to the human (hg38) transcriptomes using CLC Genomics Workbench v.23.0. Raw read counts were analyzed using the DESeq2 package implemented in R (31) using default settings to identify transcripts of significantly different abundance. These data were also used to calculate relative log2-fold change values across cell type and infection parameter, and these data were fed into pre-ranked Gene Set Enrichment Analysis (GSEA) to identify host gene sets that were negatively or positively enriched. One experiment with three replicates was performed. Fastq files have been deposited in the NCBI Short Read Archive (accession numbers pending).

### Luminex assay

To evaluate the cytokine and chemokine profile secreted by TS^CYT^ and TS^SYN^ infected or mock infected with *T. gondii*, supernatants were collected from those cells Luminex assay was performed using the following kits according to the manufacturer’s instructions: Bio-Plex Pro Human Inflammation Panel 1, 37-Plex kit (171AL001M; Bio-Rad) Bio-Plex Human Chemokine Panel, 40-plex (171AK99MR2; Bio-Rad).

In order to verify the CCL22 production in response to the infection with *T. gondii* in a different system to compare to our data with TS cells, we performed an experiment using trophoblast organoids (line TO74) (32). TOs were cultured as described previously (32), and for infection, they were removed from the Matrigel “dome” and maintained in suspension. Then, TOs were infected with 1×10^6^ *T. gondii* tachyzoites (RH-YFP) for 24h. As a control, TOs were mock infected with the parasite. The supernatants from infected and control conditions were collected and the levels of CCL22 were measured by Luminex (40-plex 171AK99MR2; Bio-Rad).

### Transmission and Scanning microscopy

For transmission electron microscopy (TEM), TS^CYTs^ and TS^SYNs^ were infected or not with *T. gondii* RH: YFP for 24h or *L. monocytogenes* wildtype for 8 h. TS^CYTs^ were fixed with 2.5% glutaraldehyde in PBS for 1 hour at room temperature and washed once with PBS. Meanwhile, TS^SYNs^ were pelleted, and the samples were kept into 2.5% glutaraldehyde. They were processed by the Center of Biological Imagining-CBI at the University of Pittsburgh. Briefly, the samples were incubated for 1 hour 4°C in 1% OsO_4_ with 1% potassium ferricyanide and washed three times with 1× PBS. Then, they were dehydrated in a graded series of alcohol for ten minutes with three changes in 100% ethanol for 15 minutes and changed three times in epon for 1 hour each. Following the removal of epon, samples were covered with resin and polymerized at 37°C overnight and then 48 hours at 60°C. After the samples were cross sectioned, they were imaged using the JEOL 1400-Plus microscope.

For scanning electron microscopy (SEM), TS^CYTs^ and TS^SYNs^ were infected with *T. gondii* for 2 h, washed twice with PBS, and the samples were fixed as described above and processed by CBI facility. Images were taken using the JSM-6335F microscope.

### Statistical Analysis

Statistical analyses (besides those on RNAseq data which were described above) were performed using GraphPad Prism 9.0 (La Jolla, CA, USA). Differences between conditions were assessed by one-way ANOVA followed by the Bonferroni multiple comparison *post-hoc* test for parametric samples, or Kruskal Wallis followed by Dunn’s multiple comparisons for a non-parametric, by two-way ANOVA, or by *t-*test for parametric samples or Mann-Whitney for non-parametric, if comparing only two groups.

## RESULTS

### Syncytiotrophoblasts are more resistant to *T. gondii* infection than cytotrophoblasts

TS^CYT^ cells were cultured and differentiated into TS^SYN^ cells as described previously (27) and we quantified the success of differentiation using the TS^SYN^ marker SDC-1 (27) and the TS^CYT^ marker ITGA-6. We found that cells in “CYT” conditions were negative for SDC-1, as expected (**Fig. 1A** and **1B**), while ∼90% of those cells cultured in 3D SYN conditions expressed SDC-1 and therefore were likely TS^SYNs^ (**Fig. 1A** and **1B**). When TS medium was supplemented only with Y27632, this resulted in a mixed population of ∼45% TS^CYT^ and 55% TS^SYN^-like cells (**Fig. 1A** and **1B**). These data indicated that we were able to recapitulate the differentiation procedure described previously (27).

**Figure 1:**
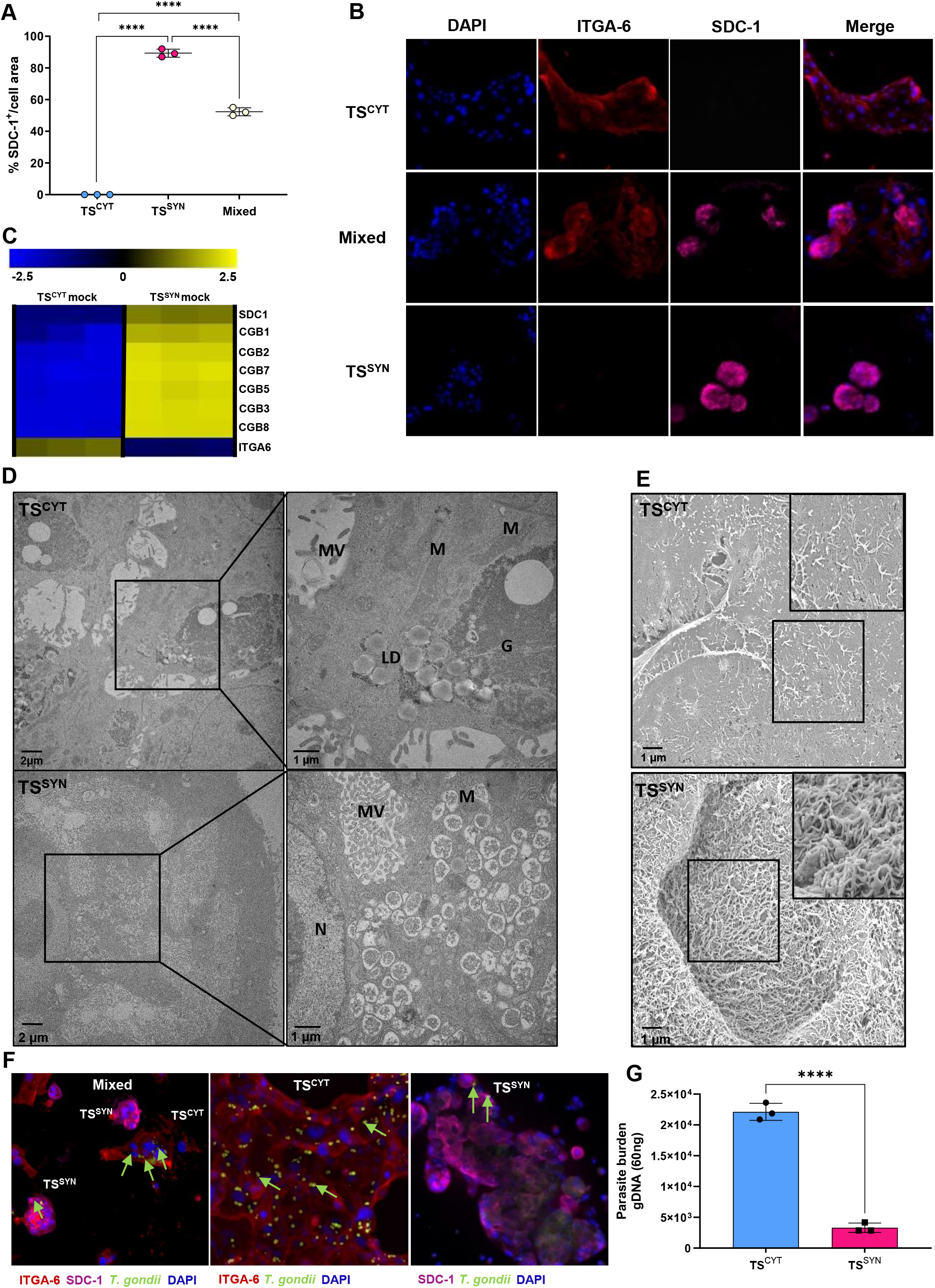
Trophoblast stem (TS) cells differentiated into syncytiotrophoblast-like cells have innate resistance to *Toxoplasma gondii*. TS^CYT^, TS^SYN^ and mixed populations of TS^SYNs^ and TS^CYTs^ were cultured from1.5 x 10^5^ cells in a 6-well plate in triplicate. Cells were infected or not with *T. gondii* with MOI of 5 for 24h prior to immunofluorescence, qPCR or RNA-seq analysis. Immunofluorescence was performed to stain cells with TS^SYN^ marker (SDC-1) (pink), TS^CYT^ marker (ITGA-6) (red), DAPI (blue) and *T. gondii* (green). (**A**) Quantification of SDC-1 in TS^CYT^, TS^SYN^ and mixed populations of TS^SYNs^ and TS^CYTs^ were calculated by the percentage (%) of fluorescence expression of SDC-1 normalized by total cell area. Differences between cells were analyzed by one-way ANOVA with Bonferroni multiple comparison *post-hoc* test. \*\*\*\**P* < 0.0001. (**B**) Representative images showing immunofluorescence microscopy of TS^CYT^, mixed populations of TS^SYN^ and TS^CYT^, and 3D TS^SYN^ (which exclusively form SYNs). Images such as these were used to generate the data in panel A. (**C**) Heat map showing transcript abundance in TS^CYT^ and TS^SYN^ for ITGA-6, CGB1, 2, 7, 3, 5, 8 and SDC-1 in mock treated and infected cells (*Padj* 0.01; log fold change ≥ 2 or ≤ −2 for all genes shown). This data provide transcriptional evidence for the establishment of TS^CYT^ and TS^SYN^ cultures in our laboratory, consistent with prior work (Okae *et al.*, 2018). (**D**) TEM photos showing the ultrastructure TS^CYTs^, highlighting the presence of lipid droplets (LD), glycogen granules (G), microvilli (MV) and mitochondria (M) and in TS^SYNs^ we observe of presence of lots of mitophagy vesicles, nuclei (N), mitochondria (MV) and a dense microvilli (MV). (**E**) SEM showing the difference in the density of microvilli in TS^CYTs^ and TS^SYNs^. (**F**) Immunofluorescence images demonstrating differences in *T. gondii* (RH: YFP) (green) infection and proliferation between TS^CYT^ and TS^SYN^ in mixed population, TS^CYT^ culture and TS^SYN^ 3D condition. Green arrows indicate parasites that are either inside or associated with the outside of host cells. (**E**) Quantifying *T. gondii* burden using qPCR for the GRA1 transcript utilizing the standard curve. Differences between TS^CYT^ and TS^SYN^ were analyzed by *t*-test, \**P* = 0.001. Scale bar: 100 μm.

To further validate the differentiation process in our laboratory we used RNAseq and first focused on transcript levels for SYN markers (CGB1, CGB2, CGB7, CGB5, CGB3, CGB8 and SDC-1) and the CYT marker ITGA-6. TS^SYN^ cells in 3D medium had significantly higher transcript abundance for SDC-1 and members of the *CGB* gene family, and significantly lower levels of ITGA-6 (**Fig. 1C**), while TS^CYT^ cells had significantly higher levels of ITGA-6 transcript and significantly lower levels of SDC-1 and *CGB* family member transcripts (**Fig. 1C**). These data suggest that we were able to generate TS^SYN^ cells and indicated that 3D medium was the most efficient protocol to generate TS^SYN^ cells as previously described (27).

We also performed TEM and SEM to evaluate differences in ultrastructure between TS^CYTs^ and TS^SYNs^. Transmission electron microscopy revealed that TS^CYTs^ have large amounts of glycogen granules and lipid droplets scattered in the cytoplasm while this is not seen in TS^SYNs^ (**Fig. 1D**). We also could observe the differences in mitochondria morphologically, similar to what described by (33). TS^CYTs^ have larger mitochondria with a lamellar crista (**Supplementary fig. 1A, B**) while TS^SYNs^ have smaller mitochondria containing vesicular cristae (**Supplementary fig.1 C-D**). Scanning electron microscopy shows that the surface of TS^SYN^ is densely covered by microvilli while microvilli in TS^CYTs^, was much more diffuse (**Fig.1E**) These findings indicate that TS^CYTs^ and TS^SYNs^ generated in our hands bear all of the expected characteristics.

Given the known resistance of SYNs to *T. gondii* infection compared to CYTs in both primary human trophoblast (PHT) cells and placental explants (18, 24), we used trophoblast stem cells to generate TS^SYNs^ and compare their sensitivity to *T. gondii* infection with TS^CYTs^. We generated pure TS^CYT^, TS^SYN^ (3D) and cultures of mixed TS^CYTs^ and TS^SYNs^ and infected them with RH: YFP parasites with an MOI of 5 for 24 h. Qualitatively, images of infected TS^CYTs^ and TS^SYNs^ show dramatic differences in *T. gondii* numbers that have infected each cell type, with TS^CYT^ cells being much more susceptible to parasite infection compared to TS^SYN^. (**Fig. 1F**). Using qPCR for the *T. gondii* gene *GRA1* as a proxy for parasite number, based on a standard curve, we found that the parasite burden was higher in TS^CYT^ cells compared to TS^SYN^ (*P* = 0.0001) (**Fig. 1E**), further supporting the reduced number of *T. gondii* associated with TS^SYN^ compared to TS^CYT^. These data suggest that TS^SYNs^ are similarly resistant to *T. gondii* as those generated during the culture of placental explants and primary human trophoblast cells (18, 24).

### TS^SYNs^ become resistant at day 4 of differentiation

To generate a pure population of TS^SYNs^ we induce the differentiation for 6 days using the ST 3D medium. Given that, we want to evaluate on which day of differentiation the cells would become resistant to *T. gondii* infection. Cells were plated in duplicate and infected on day 1, the same day they were plated, and on the following days until day 6. Cells were collected after 24 h of infection and que total of vacuoles size was calculated per cell are in a field of view. Our data shows that from day 1 to day 3 of differentiation the cells are still susceptible to infection and the parasite is able to proliferate intracellularly. However, from day 4 the cells become resistant compared to day 1 (*P* = 0.03) (**Fig. 2A**). There is no significant difference between day 4, day 5 and day 6. Representative images show the progression of the differentiation with the SDC-1 staining and the formation of the multinucleated cells over time. We also can see the large number of *T. gondii* vacuoles in the cells in day 1 of differentiation when compared to days 4, 5 and 6 (**Fig. 2B**).

**Figure 2.**
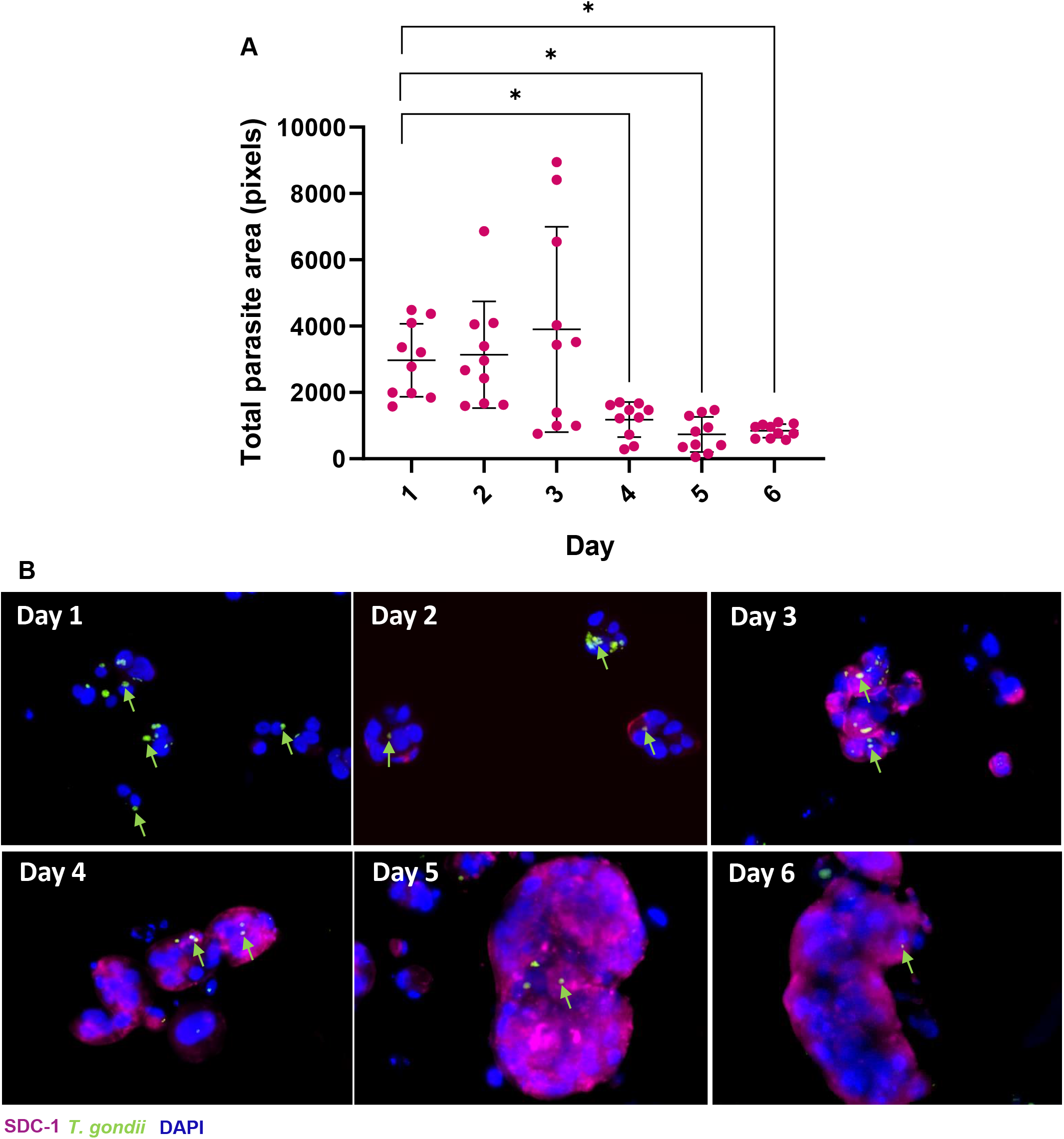
TS^SYNs^ become resistant to *T. gondii* on day 4 of differentiation. TS^SYNS^ differentiated in a ratio of 1.5 x 10^5^ in a 6-well plate in duplicate. Cells were infected with *T. gondii* every day from day 1 to day 6 and collected after 24h post-infection, fixed with 4% PFA and then visualized using immunofluorescence. (**A**) The total of parasite area was measured by Image J software in 10 fields of view. Differences between the days of differentiation and the total of parasite vacuole size was analyzed by one-way ANOVA with Bonferroni multiple comparison *post-hoc* test. (**B**) Immunofluorescence images illustrate the parasite growth in different days of differentiation from day 1 to day 6. SDC-1^+^ (TS^SYN^) in pink, *T. gondii* in green and DAPI in blue. Green arrows indicate parasites inside the host cells. Scale bar: 50 μm.

### *T. gondii* growth is restricted in TS^SYN^

A limitation of PHT cells is their short (2-4 days) cultivation time *in vitro*, making long-term quantification of *T. gondii* growth very challenging. Given the longer survival times of TS^SYNs^ derived from trophoblast stem cells, we used this model to assess parasite growth in TS^SYNs^ and TS^CYTs^ over a 48h period. To do this we infected TS^CYT^ and TS^SYN^ (3D) with *T. gondii* at an MOI of 1.5 and quantified parasite abundance using immunofluorescence at 24, and 48 h post-infection. As expected, *T. gondii* numbers significantly increased at each time point in TS^CYT^ cells (*P* < 0.0021) (**Fig. 3A, B**) and had significantly higher numbers of parasites compared to TS^SYNs^ at each time point (**Fig. 3A, B**). In contrast, parasite numbers did not significantly change over the course of the experiment in TS^SYNs^, suggesting that, similar to PHT cells (18) TS^SYNs^ restrict *T. gondii* replication (**Fig. 3A, B**). The TEM was performed in infected TS^CYTs^ and TS^SYNs^ with *T. gondii* to evaluate the differences in parasite growth and health at the ultrastructural level. The TEM photos demonstrate that the parasite grows normally in TS^CYTs^, showing a huge vacuole containing normally more than 6 health tachyzoites inside (**Fig. 4A, B**). The parasites present the normal tachyzoites morphology with characteristics organelles, but in TS^SYNs^, the TEM pictures reveal unhealthy parasites, showing few tachyzoites per vacuole and malformation tachyzoites of *T. gondii* after 24h of infection, observed by abnormal morphology, seems to be dead parasites (**Fig. 4C, D**).

**Figure 3.**
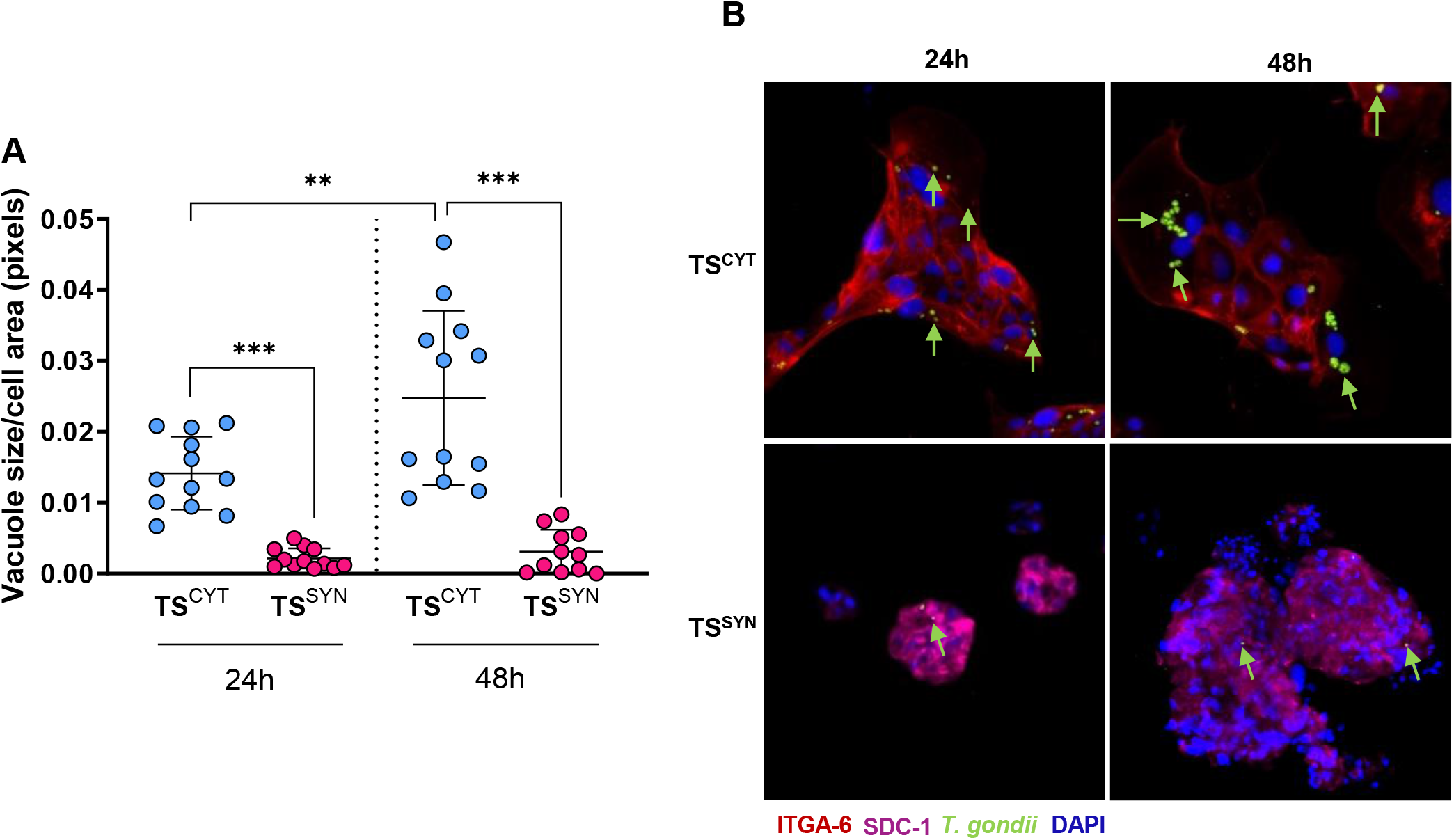
*T. gondii* growth is restricted in TS^SYN^ compared to TS^CYT^. TS^CYT^ and TS^SYN^ cells were cultured on round coverslips in 24-well plates and infected on days 3 and 5 post-plating, respectively, at an MOI of 1.5 for 24 and 48. Cells were fixed with 4% PFA and then visualized using immunofluorescence. (**A**) *T. gondii* burden in TS^CYT^ and TS^SYN^ was measured based on average vacuole size divided by total host cell area at each time point. Four images were taken from each slide using epifluorescence microscopy and analyzed using ImageJ software. (**B**) Immunofluorescence images illustrate the parasite growth in the distinct cell cultures at 24 and 48 post-infection. ITGA-6^+^ cells (TS^CYT^) are shown in red, SDC-1^+^ (TS^SYN^) in pink, *T. gondii* in green and DAPI in blue. Green arrows indicate parasites that are either inside or associated with the outside of host cells. Even after 48 h there were very few parasites associated with the TS^SYNs^. Differences between TS^CYT^ and TS^SYN^ at different time points were analyzed by one-way ANOVA with a Bonferroni multiple comparison *post-hoc* test. \**P* < 0.0001 when comparing TS^SYN^ and TS^CYT^ at each time point and when comparing across time points in TS^CYT^. Two experiments were performed with three replicates. Scale bar: 100 μm.

**Figure 4.**
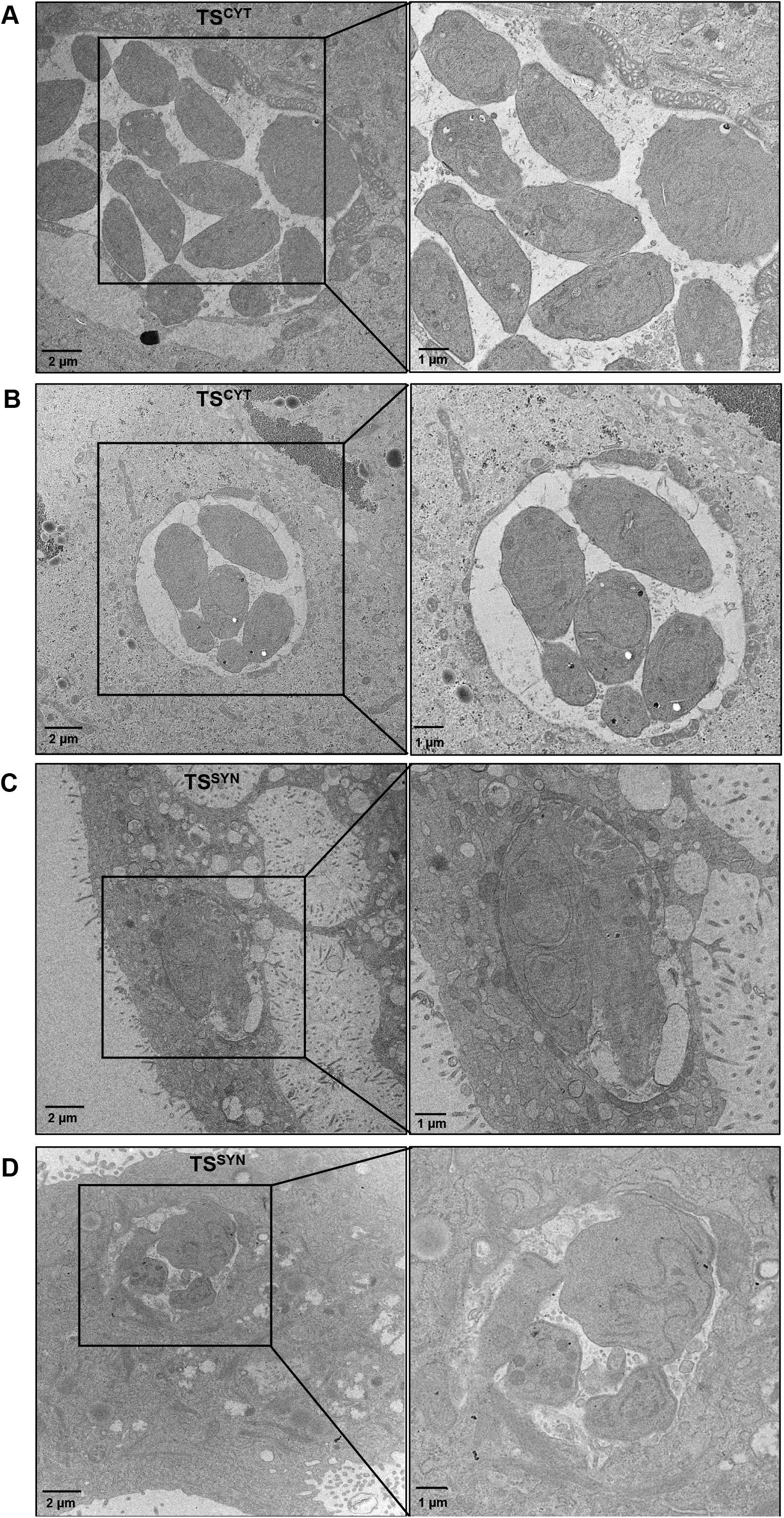
Transmission electron microscopy shows that *T. gondii* growth is restricted in TS^SYN^ compared to TS^CYT^. The TS^SYNs^ cells and TS^CYTs^ were infected with *T. gondii* for 24 h and processed for TEM. (**A-B**) shows the *T. gondii* vacuole with healthy parasites growing in TS^CYTs^. (**C-D**) *T. gondii*-containing vacuoles within TS^SYNs^ were typically smaller and often harbored parasites showing signs of low viability such as vacuolation and loss of arc-like shape. Mag: 8,000 X and 12,000 X, scale bar: 2 μm or 1 μm, respectively.

### TS^SYNs^ recapitulate the SYN-specific resistance of primary cells to *T. gondii* invasion

Since our prior work also established that SYNs from primary human trophoblasts were poorly invaded by *T. gondii*, we counted the total number of vacuoles present in each cell per field of view after 24 h post infection as a proxy of invasion rate in TS^CYTs^ and TS^SYNs^. A significantly higher number of parasite vacuoles were found in TS^CYT^ cells compared to TS^SYN^ cells per field of view (*P* = 0.0004) (**Fig. 5A**), indicating that *T. gondii* invaded more TS^CYTs^ compared to TS^SYNs^. This phenotype recapitulates what was observed previously in PHT cells (18). Specifically, fewer parasites invaded TS^SYNs^ compared to _TSCYTs._

**Figure 5.**
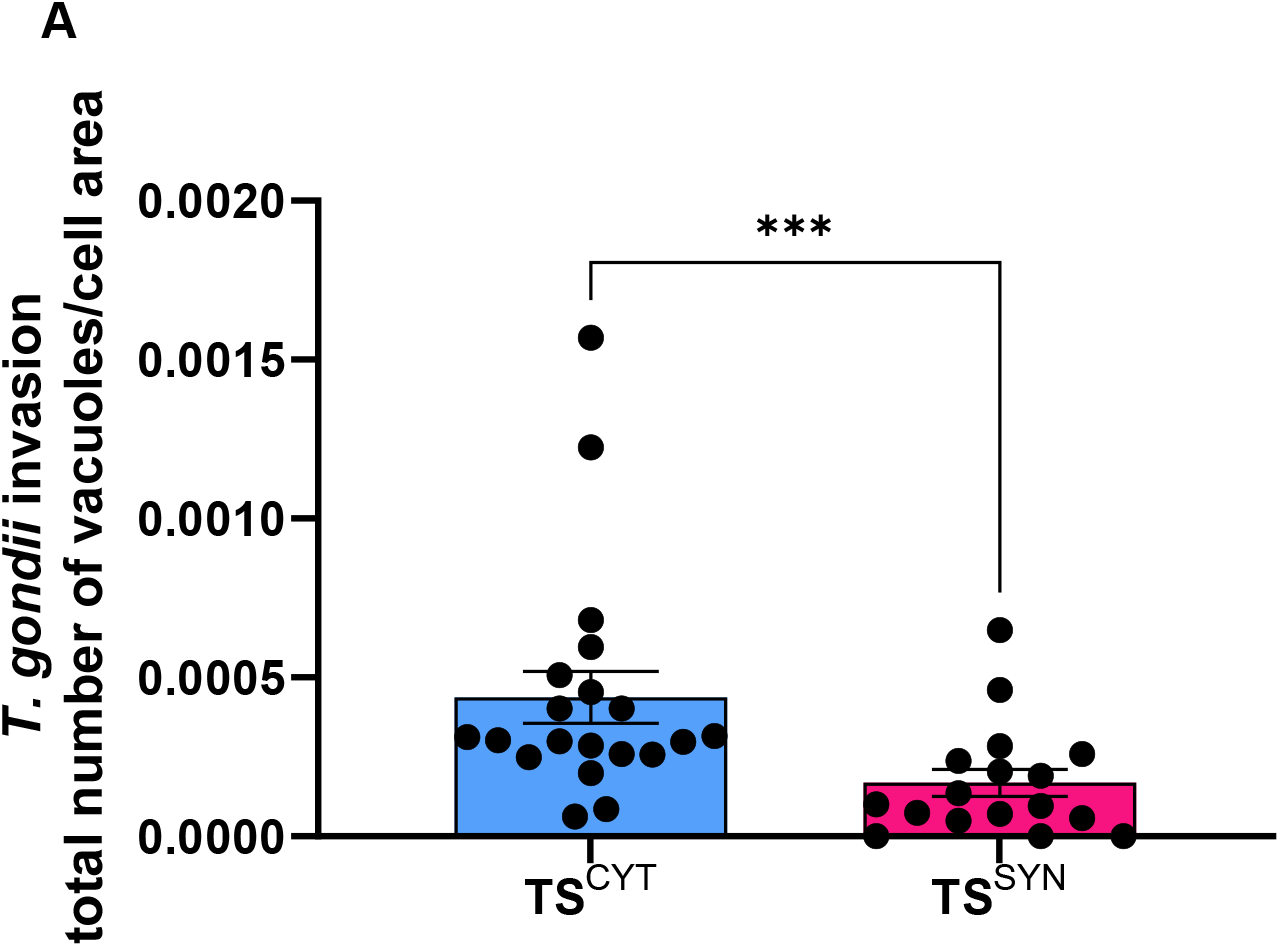
TS^SYN^ are less susceptible to *T. gondii* invasion compared to TS^CYT^. TS^CYT^ and TS^SYN^ were cultured on glass slides in 24-well plates and infected on days 3 and 5 post-culture, respectively, with an MOI of 5 for 24h. Cells were fixed with 4% PFA and the immunofluorescence was performed. (**A**) Quantification of *T. gondii* invasion in TS^CYT^ and TS^SYN^ was calculated by the total number of vacuoles with greater than one parasite per total cell area per field of view. (Differences between TS^CYT^ and TS^SYN^ were calculated by Mann-Whitney, \**P* = 0.0004). Two independent experiments with three replicates were performed.

### TS^CYTs^ and TS^SYNs^ exhibit low production of immunoregulatory factors compared to PHTs

In addition to differences in susceptibility to *T. gondii* compared to many other cell types, PHTs, human placental villous explants and trophoblast organoids produce different cytokines, chemokines and grow factors that are important for pregnancy maintenance and fetus defense (2, 32). Previous work has shown that CCL22 is produced by PHTs and villi in response to *T. gondii* infection while other cell types such as HFFs do not (18). We quantified 77 chemokines and cytokines using multianalyte Luminex-based profiling in supernatants of mock- and infected cells to evaluate the immunomodulatory secretome of TS^CYTs^ and TS^SYNs^. Even though TS cells recapitulate the resistance profile to *T. gondii* infection, the cells do not recapitulate the immune response to infection observed in villous explants and PHT cells. In our data we defined as reliably secretion factors measured above 100 pg/mL as baseline. We found that both TS^CYTs^ and TS^SYNs^ did not secrete CCL22 when infected with *T. gondii* (**Fig. 7A, B** and **C**). In contrast, when we infected trophoblast organoids (TOs), we detected a clear induction of CCL22 secretion compared to mock-infected organoids (**Fig. 7D**). Our data also show that TS^CYTs^ in both secrete cytokines as (MIF, IL-11 and gp130), factors (TNFRSF8, Pentraxin-3, TNFRSF8 and MMP-1), and two soluble factors (TNF-R1 and OPN) (Fig.7A). In other hand, TS^SYNs^ only secrete detectable amounts of two soluble factors (OPN and TNF-R2) and one cytokine (MIF) (**Fig.7B**). Interestingly, we observed that the soluble factor OPN presents a significant low levels of secretion in TS^SYNS^ after infection when compared with TS^CYTs^ infected **(Fig. 7A, B** and **D**), and the cytokine MIF was highly produced by TS^CYTs^ and TS^SYNs^ in both conditions (**Fig. 7A, B** and **F**), however the infection with *T. gondii* induce significantly higher levels of this cytokine compared to the mock conditions, and infected TS^SYNs^ also release more MIF when compared to TS^CYTs^ (**Fig. 7A, B and F**).

**Figure 6.**
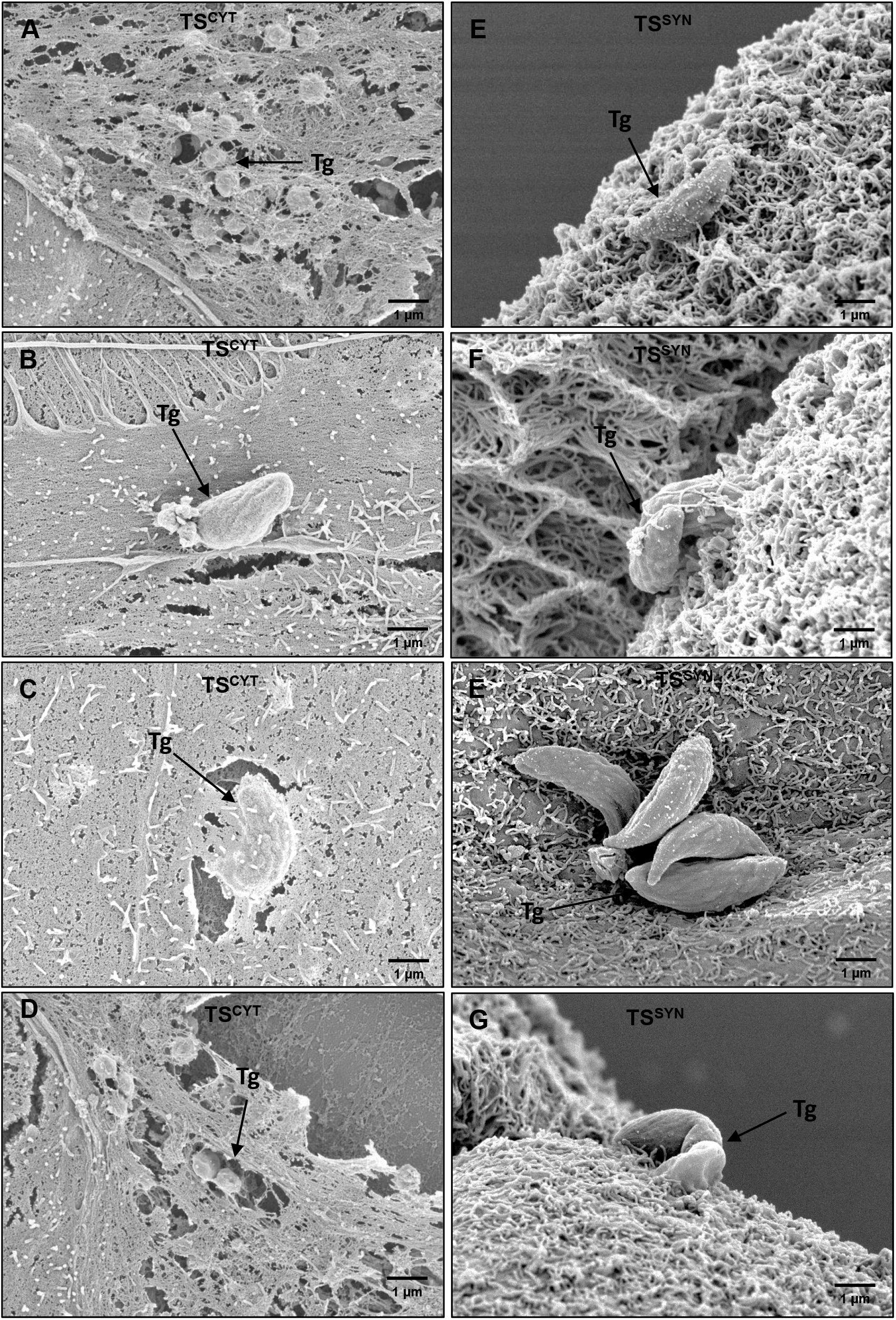
Scanning electron microscopy in TS^CYTs^ and TS^SYN^ infected with *T. gondii*. TS^CYT^ and TS^SYN^ were infected for 2h and samples were collected for SEM and pictures were taken using the SEM microscopy. (**A-D**) qualitative images showing the parasite invasion process in TS^CYTs^, where we can visualize a large number of parasites under the membrane, suggesting successful invasion process. (**E-G**) qualitative images show parasites associated with TS^SYNs^. *Toxoplasma gondii=* Tg. Black arrows indicate the parasite. Mag: 10,000 X, scale bar: 1μm.

**Figure 7.**
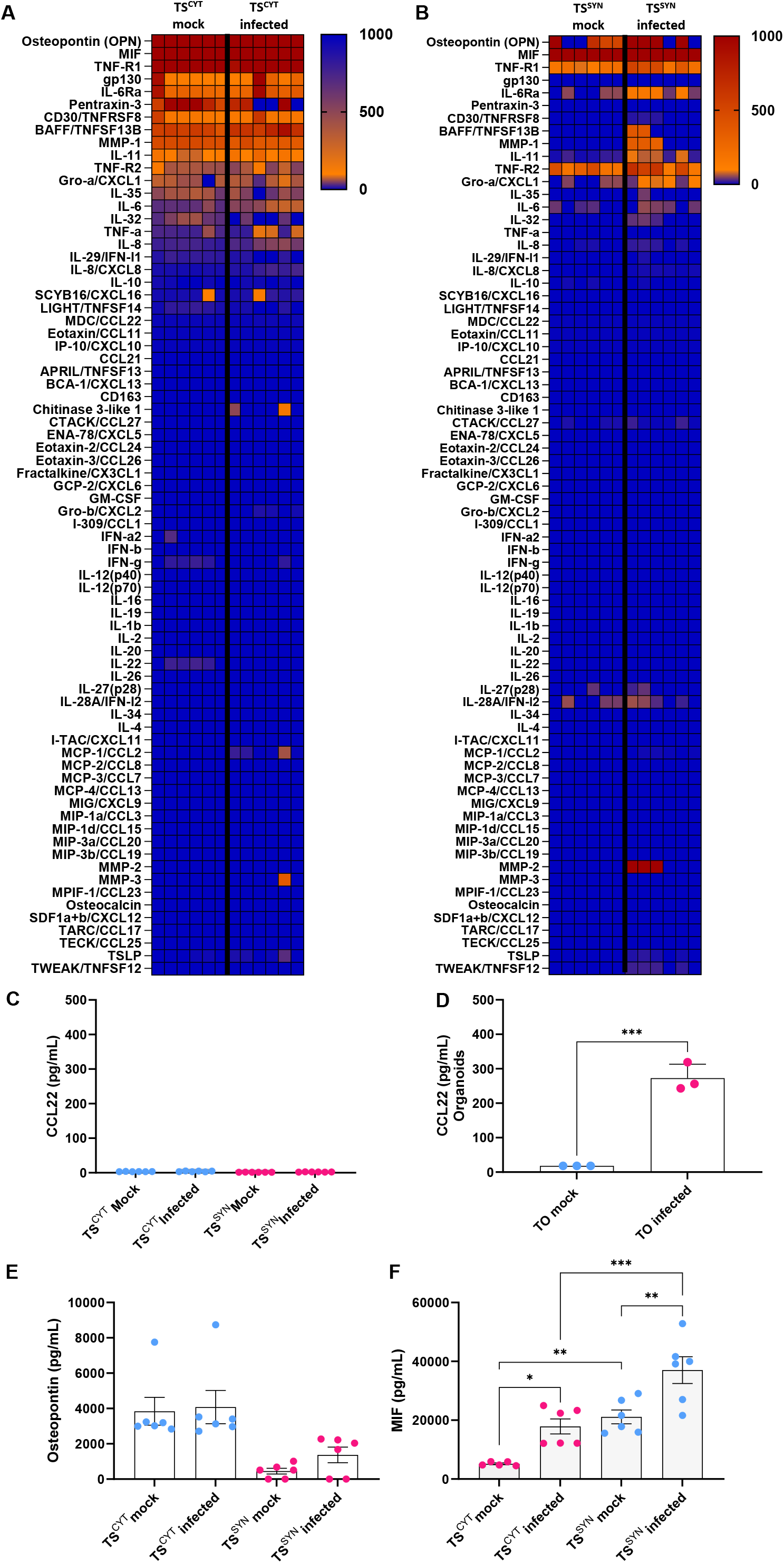
Cytokine quantification in supernatants from TS^SYN^ and TS^CYT^ and trophoblast organoids mock infected or infected with *T. gondii.* The supernatants of TS^CYT^, TS^SYN^ and TOs of mock or infected with *T. gondii* were collected after 24 h post-infection and the Luminex assay was performed to visualize the immunomodulatory profile in both cells and the induction of CCL22 of TOs. The heatmap graphs show the difference in secretome levels of different cytokines, chemokines, and immune factors in (**A**) TS^CYT^ mock or infected and (**B**) TS^SYN^ mock or infected with *T. gondii*. Differences among the secretion of CCL22 (**C**) in both cell types mock and infected and in (**D**) TOs. We also highlight the differences in secretion levels of (**E**) osteopontin and (**F**) MIF in TS^CYTs^ and TS^SYNs^ mock and infected with *T. gondii.* The data was expressed in pg/mL. Differences between TS^CYT^ and TS^SYN^ at the cytokine level were analyzed by one-way ANOVA with a Bonferroni multiple comparison *post-hoc* test.

Overall, our data show that trophoblast stem cells do not release many of the immunomodulatory compounds that are secreted by PHTs, highlighting that these cells do not recapitulate the similar immunological profile of PHTs and placenta explants. This model, then, allows us to separate immune effector and signaling molecule production from innate cellular resistance that develops during the transition from CYT to SYN.

### *T. gondii-*infected TS^SYN^ and TS^CYT^ have distinct transcriptomes

Another disadvantage of the PHT cell model is that the cultures are a mixed population of CYTs, SYNs, and also contain contaminating fibroblasts (18, 21, 34). The TS model described here provided a unique opportunity to observe the transcriptional response to infection in pure populations of cells that bear biological similarity to naturally occurring CYTs and SYNs (18, 27). After infecting TS^CYT^ and TS^SYN^ for 24 h with *T. gondii* RH: YFP, we performed strand specific RNAseq to compare the transcriptional responses of each of these cell types. We first used principal component analysis (PCA) to broadly examine sample-by-sample differences in the presence and absence of *T. gondii.* Two major principal components were identified. PC1 encompassed 96% of the total variance with cell type (TS^SYN^ or TS^CYT^) varying primarily along this axis (**Fig. 8A**). The bulk of the remaining variance (3% out of 4%) was accounted for by infection state (mock or infected; **Fig. 8A**). These data confirm that TS^CYTs^ and TS^SYNs^ are transcriptionally distinct, as expected from prior transcriptional studies on these cell types (27). One surprise in these data was that the impact of infection on the TS^CYT^ and TS^SYN^ transcriptomes was similar between these cell types, despite the dramatic difference in parasite infectivity (e.g., **Fig. 1F** and **1G**; **Fig. 3A** and **3B**). While we explore this further below by examining the specific sets of genes with altered transcript levels under each condition, this was a surprising result given that many of the well-known alterations in the host transcriptional profile requires parasite invasion and/or parasite attachment to the host cell along with the secretion of host modulatory effectors (35, 36).

**Figure 8.**
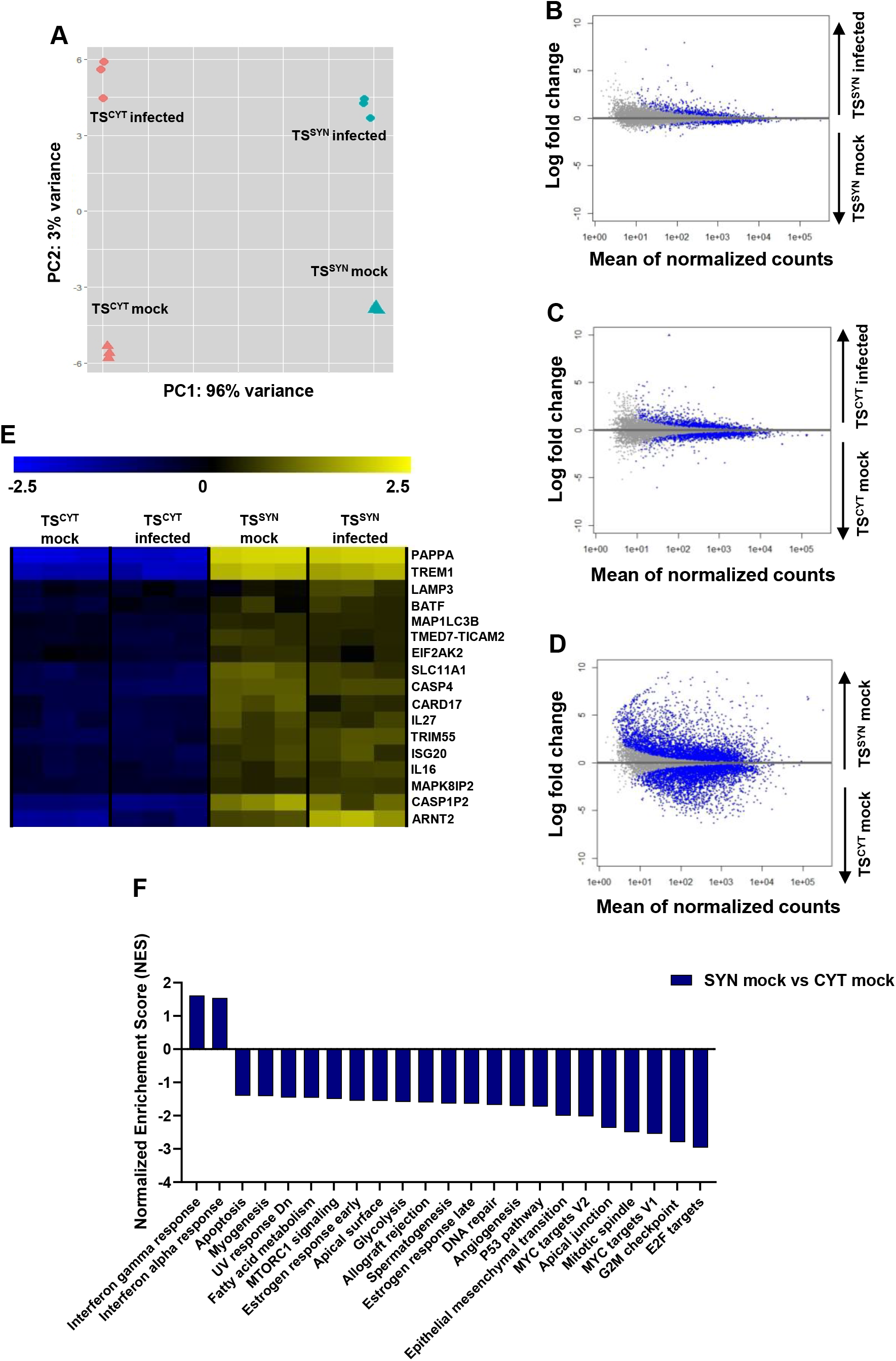
TS^SYN^ and TS^CYT^ infected cells reveal distinct gene expression profiles. Cells cultured in 6-well plates, and mock treated or infected with *T. gondii* on days 3 and 6 post plating, respectively, with an MOI of 5 for 24h. Cells were collected and processed for RNA-sequencing. (**A**) Principal components PC1 and PC2 of TS^CYT^ infected or mock infected and TS^SYN^ infected or mock infected differentiated samples along the cell type (PC1) or infection status (PC2) axes. (**B-D**) MA-plots of transcript abundance in TS^CYT^ and TS^SYN^ mock treated or infected with *T. gondii*. Blue dots represent genes of significantly different abundance based on the statistical comparison being performed: (**B)** TS^SYN^ infected versus TS^SYN^ mock; (**C**) TS^CYT^ infected versus TS^CYT^ mock; (**D**) TS^SYN^ mock versus TS^CYT^ mock. These plots illustrate that the most dramatic differences between the samples are driven by the cell type (TS^CYT^ or TS^SYN^). (**E**) Heat map showing immunity-related transcripts that were either constitutively different between cell types independent of infection status or altered in abundance by infection. (*Padj <* 0.05; log fold change ≥ 1 or ≤ −1). We used normalized enrichment scores (NES) generated using Pre-ranked Gene Set Enrichment Analysis (GSEA) from rlog-normalized data to evaluate the differences in enriched pathways between TS^SYN^ and TS^CYT^. (**F**) GSEA plot shows different gene set pathways related to metabolism in TS^SYN^ mock vs TS^CYT^ mock. For the graph, only significantly enriched pathways are shown (FDR q-value < 0.05). One experiment with three replicates was performed.

All transcripts and those of significantly different abundance based on DESeq2 analysis (Padj < 0.05; |log_2_fold-difference| > 2) are shown in the MA-plots (log fold-difference vs average abundance for each transcript) in **Fig. 8B-8D**. As shown in **Fig. 8B** and **8C**, MA plots comparing infection in both TS^SYN^ and TS^CYT^ have similar shapes and profiles, and but more significantly different transcript abundances are seen in in TS^CYTs^ after infection, compared to TS^SYNs^. A unique feature of the SYN is its ability to resist *T. gondii* infection (18, 24)without prior exposure to activating cytokines like interferon γ. Given that TS^SYNs^ recapitulate this phenotype, we aimed to identify putative host resistance genes that might be either constitutively expressed in TS^SYNs^ compared to TS^CYTs^ and other susceptible cell types or induced by *T. gondii* infection uniquely in TS^SYNs^. Our data show that some genes involved in proinflammatory or autophagic response are of significantly greater abundance in TS^SYN^ compared to TS^CYT^ in both mock-treated and infected cells, including *PAPPA, CARD17, TREM1, TMED7-TICAM2*, *BATF, SLC11A1, IL27, ISG20, MAK8 IP2, MAP1LC3B, LAMP3, IL16, CAPSP1P2, CASP4, TRIM55, ARNT* and *EIF2AK2* (**Fig. 8E**).

Using GSEA on our RNAseq dataset comparing TS^SYN^ mock versus TS^CYT^ mock identified 23 “Hallmark” pathway gene sets (FDR-q value < 0.01) (**Fig. 8F**). Interestingly, the pathways *IFNα-* and *IFNγ-response* were both significantly enriched in uninfected (“mock”) TS^SYN^ compared to uninfected TS^CYT^. All the other significant pathways are significantly downregulated in TS^SYNs^ compared to TS^CYTs^, for example, *fatty acid metabolism, MTORC1 signaling, apical surface, glycolysis, P53 pathway, epithelial mesenchymal transition, MYC target V1 and V2, apical junction and E2F targets* (**Fig. 8F**). This is consistent with a distinct response of these cell types to infection as would be expected given the dramatic differences in their transcriptional profiles (**Fig. 8A** and **8D**).

### TS^SYNs^ recapitulate resistance to *Listeria monocytogenes* infection compared to TS^CYTs^

In effort to characterize how TS^SYN^ resist infection of other teratogens besides *T. gondii*, we tested susceptibility of the cells to the congenital pathogen *L. monocytogenes* (*Lm*). When infected with WT *Lm*, far fewer CFUs are recovered from TS^SYN^ compared to TS^CYT^ (**Fig. 9A**). We observed the same deficit in recovered CFUs when TS^SYN^ and TS^CYT^ infected with *Lm* Δ*prsA2*. PrsA2 is a secreted protein chaperone that is required for the activity of several secreted *Lm* virulence factors and, here, produces an intermediate phenotype (**Fig. 9A**) (37, 38). Finally, there is no significant difference in CFUs recovered from TS^SYNs^ compared to TS^CYTs^ infected with *Lm* Δ*hly* (encodes listeriolysin O, LLO). LLO is a pore-forming toxin that is required for bacterial escape from the host cell vacuole which allows for intracellular growth and infection of neighboring cells (**Fig. 8A**)(39). The use of three *Lm* genotypes here depicts a spectrum of permissiveness in TS^CYTs^ whereas TS^SYNs^ entirely restrict *Lm* infection and persistence no matter what *Lm* genotype.

**Figure 9:**
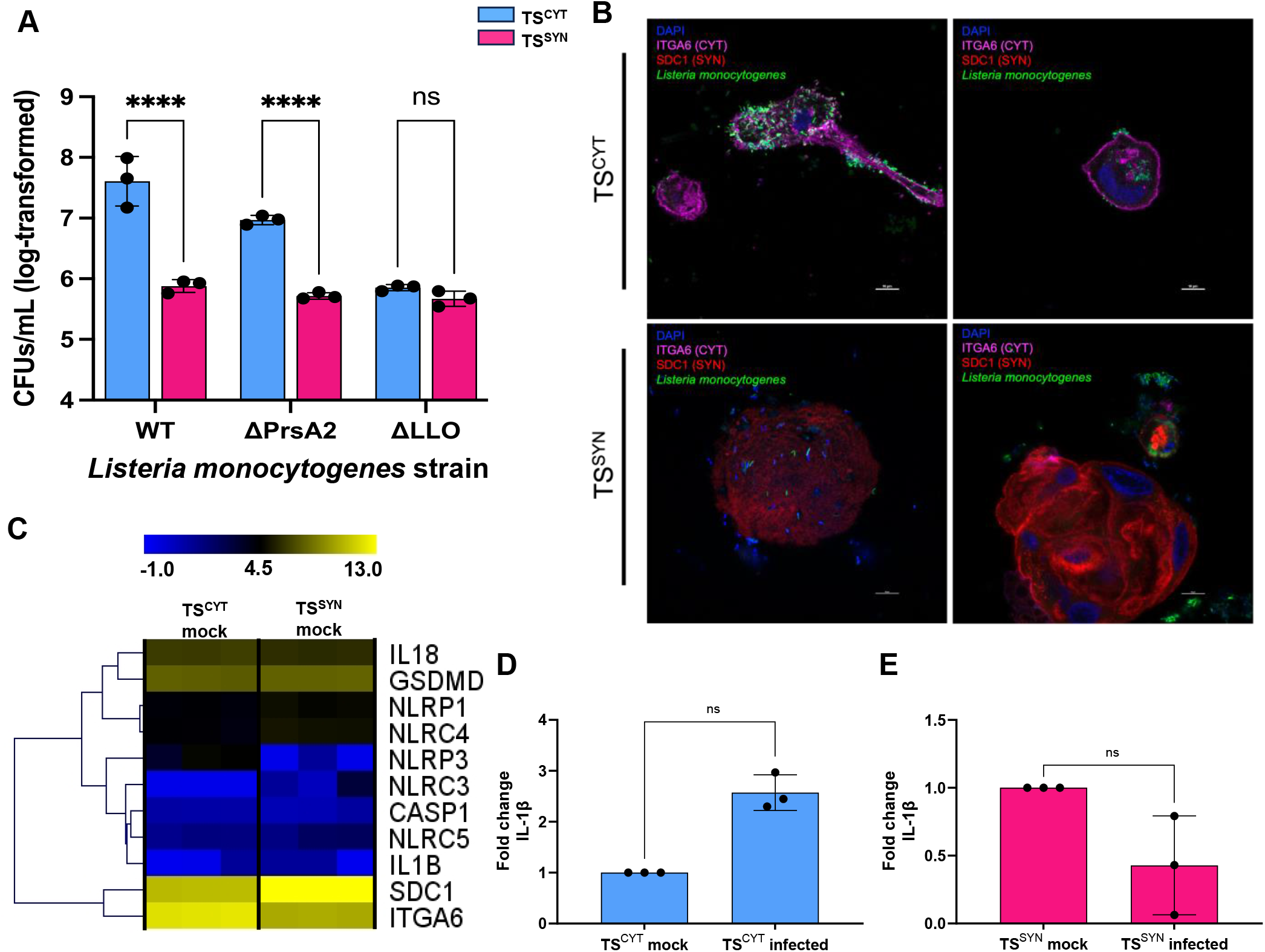
TS^SYNs^ are resistant to *Listeria monocytogenes* compared to TS^CYTs^. Cells were infected with wildtype (10403S) strain and isogenic ΔprsA2 (NF-L1651), Δhly (DP-L2161) of *L. monocytogenes* for 8 h. (A) Colony forming units (CFUs) detected on BHI agar plates after 8 hours of Lm infection in TS^CYT^ and TS^SYN^. All data is log-transformed for visualization and analysis. For all, * = p < 0.05, ** = p < 0.01, *** = p < 0.001, **** = p < 0.0001, ns = not significant. Data are analyzed with a two-way ANOVA with Holm-Šidák multiple comparisons test. Lm strain, cell type, and interaction are all *P* < 0.0001. Only pre-planned comparisons were made to minimize Type I error and those are shown on the graph. (**B**) Representative confocal images of GFP-tagged Lm-infected TS^CYT^ or TS^SYN^ for 8 hours. ITGA6 (TS^CYT^), SDC1 (TS^SYN^), DAPI (nuclei) and GFP (Lm). Two experiments in three replicates were performed. (**C**) Heat map showing transcript abundance of uninfected TS^CYT^ and TS^SYN^ for genes involved in the inflammasome pathway to illustrate relatively lower expression of these transcripts compared ITGA6 and SDC1 as representative markers CYTs and SYNs. (**D**) Quantification of the abundance abundance of IL-1β transcript using qPCR normalizing to β-Actin as a control. Differences between TS^CYT^ mock and TS^CYT^ infected, and TS^SYN^ mock and TS^SYN^ infected were analyzed by *t-*test showing no significant induction of either transcript in Lm-exposed TS^SYNs^ or TS^CYTs^. Two experiments with three replicates were performed.

Confocal imaging of GFP-tagged *Lm* infecting either TS^CYT^ or TS^SYN^ (**Fig. 9B)** mirrors the differences in recovered CFUs depicted for the WT strain (**Fig. 9A)**. We see more *Lm* attached and internalized by TS^CYT^ compared to TS^SYN^. The *Lm* located in TS^SYN^ appears primarily associated with the cell membrane but not intracellular. This distinction is seen clearly in **supplementary video 1** as there are no visible intracellular *Lm* in the 3D reconstruction of TS^SYN^ infected with GFP-tagged *Lm*. In TS^CYT^ there are more GFP-tagged *Lm* both associated with the membrane and intracellular. This is visible in **supplementary video 2** of TS^CYT^ infected with GFP-tagged *Lm*, especially in contrast to TS^SYN^.

Previous work has shown that placental trophoblast constitutively secretes the inflammasome cytokines as IL-1β and IL-18, and the infection with *L. monocytogenes* can also induces more activation of this pathway, leading to resistance against the bacteria infection (23). Due to that, we evaluate the gene expression level of some important constituents of the inflammasome pathway, and we identified those genes are expressed in low abundance in both cells TS^CYTs^ and TS^SYNs^ (**Fig. 9C**). We also measured the gene expression level of IL-1β in infected cells with *L. monocytogenes*, and the infection in both cells does not induce the gene expression of this cytokine when compared to the respective mock conditions (**Fig. 9D, E**).

We also performed TEM in TS^CYTs^ and TS^SYNs^ infected with *Lm*. The representative EM photos confirm the data found in the confocal imaging (**Fig. 9B**), in which most of the *Lm* was associated to the cell membrane of TS^SYNs^; however, in TS^CYTs^ *Lm* are successfully internalized and grow intracellularly (**Fig. 10A-10D**).

**Figure 10.**
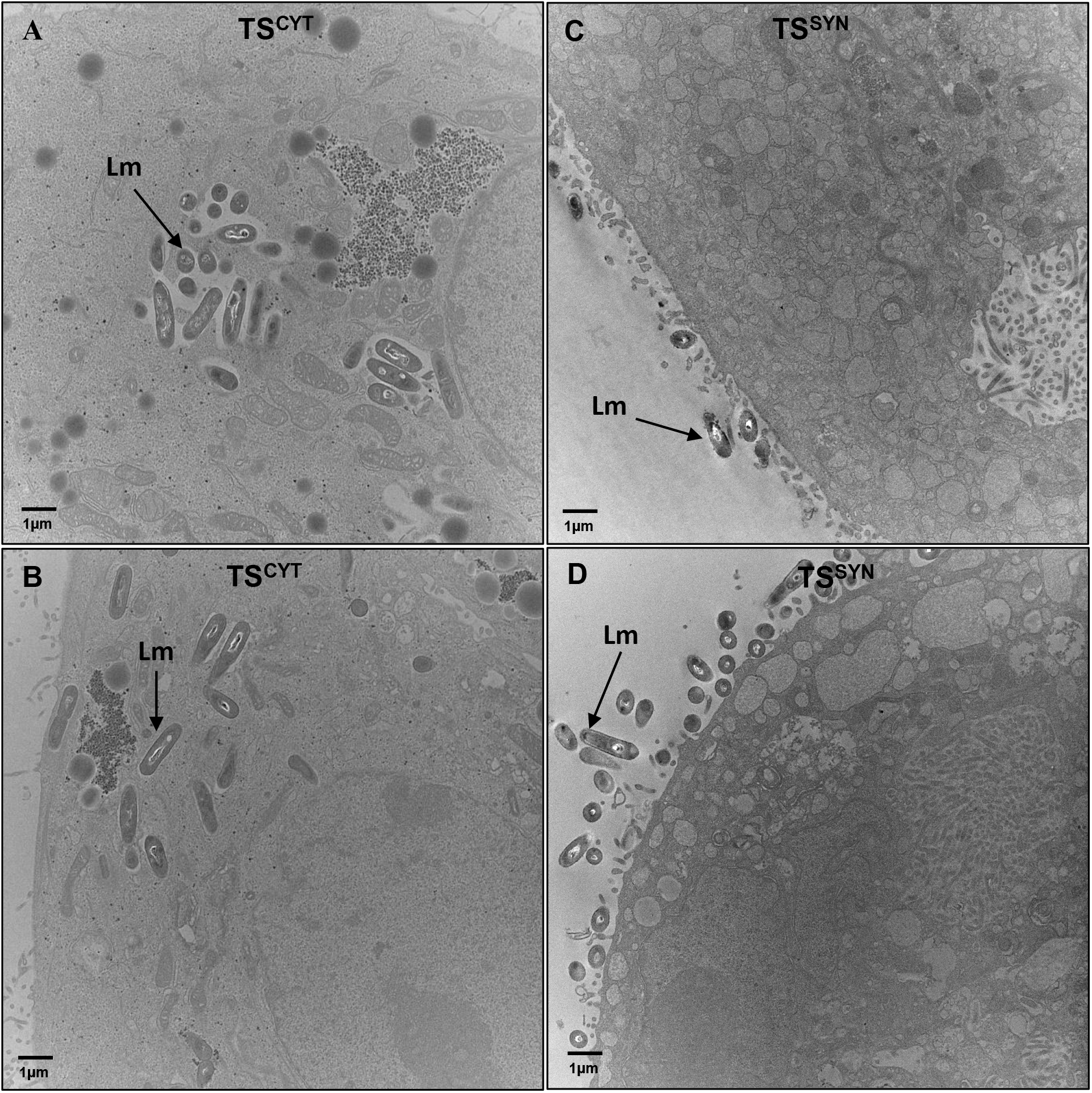
Transmission electron microscopy in infected TS^CYTs^ and TS^SYNs^ with *L. monocytogenes.* TS^CYTs^ and TS^SYNs^ were infected with *L. monocytogenes* (WT) for 8h and processed for TEM. (**A** and **B**) show the intracellular *L. monocytogenes* in TS^CYTs^. (**C** and **D**) show *L. monocytogenes* associated with the TS^SYNs^ membrane. Lm: *Listeria monocytogenes.* Black arrows indicate the bacteria. Mag: 10,000 X, scale bar: 1 μm.

## DISCUSSION

Here we describe a new *in vitro* system developed (27) to study placenta-pathogen interactions evaluating the differential susceptibility of TS^SYNs^ and TS^CYTs^ against *T. gondii*. The architecture of placenta villous explants, comprised primarily of trophoblast cells, is associated with safeguarding of the fetus against potential maternal bloodborne microbes, but the mechanisms underlying the resistance profile is still poorly understood (2, 16).

Nearly all mammalian cells studied to date can be infected with *T. gondii* and support rapid *T. gondii* replication, unless they are made resistant by exposure to effector cytokines like interferon-γ. However placental syncytiotrophoblasts are an exception to this rule, having clear innate resistance to *T. gondii* that has been demonstrated in multiple primary placental models including villous explants and primary human trophoblast (PHT) cells (18, 24). The mechanisms involved in SYN resistance to parasite adhesion and ability to restrict parasite growth are unknown (16, 18), and a significant barrier to elucidating these mechanisms are genetically tractable and reproducible models of CYT and SYN development. Overall, our data provide evidence that TS-derived CYTs and SYNs recapitulate the susceptibility and resistance phenotypes that we and others have previously characterized. Most notably, TS^SYNs^ resist *T. gondii* infection at the level of both adhesion and replication, which is identical to what we have observed previously using mixed CYT and SYN cultures derived from term placentas (18). This same dichotomy is observed in midgestational chorionic villi (18, 40).

Additionally, another study using villous explants from the first trimester showed that SYN act as a strong barrier against *T. gondii*, since these cells were also very resistant to infection (24). Mechanisms of SYN resistance to a variety of pathogens seems to depend on the infection model. Our data showed that besides the resistance against *T. gondii* infection, TS^SYNs^ also were resistant to *L. monocytogenes,* mainly by restricting the bacterial entry into the cells. Looking at *L. monocytogenes* and *T. gondii* infection, we can observe that TS^SYNs^ are very resistant to pathogen invasion/entry, similar to what is shown in SYNs from PHTs and villous explants (18, 22, 41). SYNs lack intracellular junctions, which some bacterial and viral pathogens use to invade cells (42, 43). They also have a robust cytoskeletal network and branched microvilli that might inhibit pathogen entry, and in fact are highly resistant to *T. gondii* adhesion, a required first step for *T. gondii* to ultimately infect a mammalian cell (18). Our work here shows a clear difference in the density of the microvilli between TS^SYNs^ and TS^CYTs^, in which the cell surface of TS^SYNs^ is highly density covered by microvilli while in TS^CYTs^ is not. SEM photos from both infected cells for 2 h clearly show more parasites that invade TS^CYTs^ (**Fig. 6A-6D**), and in TS^SYNs^, *T. gondii* seems to be attached to the membrane, but rarely invades the cells in the same time frame as compared to TS^CYTs^ (**Fig. 6E-6G**).

Interestingly, besides the presence of microvilli, TS^SYNs^ have a significant decrease in the transcriptome abundance of a large number of genes related to apical surface and apical junctions, including *ICAM1, TJP1, CDH1, CDH3*, that having described be important for pathogens invasion, sharing the same features as SYNs from PHTs cells (**Supplementary fig. 2A**). So, we suggest that these features could be involved in the differential invasion/entry rates of *T. gondii* and *L. monocytogenes* in TS^CYTs^ compared to TS^SYNs^. Surface proteoglycan content may also differ in TS^SYN^ compared to TS^CYT^, a possible mechanism supported by both RNAseq data in the present study and others (Okae *et al.*, 2018) that TS^SYN^ have lower transcript abundance for HSPG2, and ICAM-1 compared to TS^CYT^ (**Table S1**). HSPG2 and ICAM-1 are involved in *T. gondii* attachment and invasion, being targeted by parasite surface proteins such and SAG-3 and MIC-2, respectively to promote invasion (44, 45). Host cell surface proteoglycans are also generally critical for *T. gondii* adhesion (46), and our transcriptional data also show a clear reduction in the levels of XYLT1 transcript during the TS^CYT^ to TS^SYN^ conversion *in vitro* (**Table S1**), which catalyzes one of the first steps in proteoglycan synthesis by adding xylose to serine residues in target proteins (47). This is another possible means of restricting pathogen adhesion to SYNs, in particular for those that require preliminary adhesion events to proteoglycans. Lectin-based studies have also demonstrated clear differences in surface sugar content across different placental cell types including CYTs and SYNs (48). The TS^SYN^ and TS^CYT^ system could be used to study the role of surface proteoglycan content on cell-specific restriction in *T. gondii/L. monocytogenes* adhesion given its reproducible growth and differentiation characteristics and genetic tractability.

In addition to its physical barrier function, the trophoblast triggers a powerful immune response by releasing various cytokines and immunological factors. Trophoblasts produce cytokines constitutively and in response to infection, including those associated with the inflammasome, such as IL-1β and IL-18, which control *L. monocytogenes* infection (23). In contrast, CCL22 is only detected in large quantities following infection of PHT cells with *T. gondii* (18, 28). CCL22 also increases in abundance during miscarriage (49, 50). Our data showed that both TS^CYTs^ and TS^SYNs^ do not recapitulate the immunological secretome previously observed in PHT cells and villous explants, and in the present study placental organoids (**Fig. 7D**). Therefore, with respect to *T. gondii*, at least, TS^SYNs^ can be used to directly explore the structural impediments to parasite adhesion and mechanisms of IFNγ-independent restriction of parasite replication, while other models like the placental organoid are more useful for studying both basal and induced immunological mechanisms of resistance.

The TS model permits us to circumvent one limitation of the PHT model which is that the cultures are a mixture of both CYT and SYN cells that vary in their ratios between preparations. Given that TS-derived CYTs and SYNs can be cultivated in a manner that leads to relatively pure cultures of a given cell type, we could examine putative CYT- and SYN-specific responses in isolation. We observed large differences in the transcriptional responses of each cell type, but one of the more remarkable findings was that although *T. gondii* invaded and proliferated poorly in TS^SYNs^, we still observed considerable changes in the transcriptional profile of these cells that rivaled those found in the more readily infected TS^CYTs^. The changes that we observed in the TS^SYN^ could be driven more by paracrine responses to the presence of *T. gondii* rather than infection, although this would have to be investigated directly. It is also possible that the TS^SYN^ are particularly sensitive to alterations induced by even the small number of invaded parasites, and/or that resistance pathways that drive the clear phenotype of restricting *T. gondii* replication by the TS^SYN^ are robustly activated by *T. gondii*.

Human trophoblast stem cells have emerged in recent years as an important tool in studying placental development as well as pathogen resistance and responses. Here we show that these cells recapitulate primary human trophoblast and explant resistance phenotype profiles, with TS-derived SYNs being highly resistant to *T. gondii* infection and being ultra structurally similar to primary cells. TS-derived SYNs also resisted infection with *L. monocytogenes* (a feature shared with placental explants(22, 23), suggesting that resisting pathogen adhesion/attachment may be a generalized mechanism of SYN-resistance.

However, the TS model has some limitations, most notably in its poor recapitulation of both constitutive and pathogen-induced cytokine production which is observed in primary trophoblast cultures and placental explants (18). These cells are genetically tractable tools to investigate cell-intrinsic mechanisms of resistance to pathogen adhesion and replication.

## AUTHOR CONTRIBUITION

RJS participated in design and performed experiments, analyzed the data, and wrote the manuscript. LFC helped in acquisition of data and analyzed the data, JLG helped in the acquisition of data, LAC helped supervised the experiments. HY helped in the acquisition of data. CBC helped in the acquisition of data. JPB, the principal investigator, participated in experimental design, supervised the experiments, and reviewed the manuscript.

## ACKNOLEGMENTS

We thank D. Stolz, M. Sullivan, J. Franks, and Ming Sun (CBI-University of Pittsburgh) for processing the samples and for technical assistance with TEM and SEM.

## CONFLICT OF INTERST

The authors declare that they have no conflict of interest.

---

**Supplementary figure S1. Transmission electron microscopy in TS^CYTs^ and TS^SYNs^.** TS^CYT^ and TS^SYN^ were processed for TEM to differences in cells organelles in both cells. Representative TEM images show the mitochondria in TS^CYTs^ (**A** and **B**) and in TS^SYNs^ **(C** and **D**). Mitochondria= M. Mag. 60,000 X, scale bar: 200nm.

**Supplementary figure S2. Apical surface and apical junction genes set.** Using the bulk RNAseq data generated by TS^CYTs^ and TS^SYNs^ mock and infected with *T. gondii* we used the Pre-ranked Gene Set Enrichment Analysis (GSEA) from rlog-normalized data to evaluate the differences in enriched pathways between TS^SYN^ and TS^CYT^. Among the gene sets that are significantly downregulated in TS^SYNs^ (FDR q-value < 0.05), we evaluated the apical surface and apical junction genes. (**A**) Heat map showing the apical surface and apical junction gene sets that have significantly different transcript abundance between TS^CYTs^ and TS^SYNs^.

**Supplementary video 1**. **3D reconstruction of TS^CYT^ infected with GFP-tagged *Lm.*** TS^CYTs^ were infected with GFP-tagged *Lm* for 8 hours and cells were analyzed by confocal microscopy. Images in z-stack were made to develop the 3D reconstruction. TS^CYT^ marker (ITGA-6) (pink), DAPI (nuclei) and *Lm* GFP (green).

**Supplementary video 2**. **3D reconstruction of TS^SYN^ infected with GFP-tagged *Lm.*** TS^SYNs^ were infected with GFP-tagged *Lm* for 8 hours and cells were analyzed by confocal microscopy. Images in z-stack were made to develop the 3D reconstruction. TS^SYN^ marker SDC1 (red), DAPI (nuclei) and *Lm* GFP (green).

**Supplementary table S1:** Log2 (FPKM) transcript count values from TS^SYNs^ and TS^CYTs^ mock and infected with *Toxoplasma gondii*.

